# When host populations move north, but disease moves south: counter-intuitive impacts of climate warming on disease spread

**DOI:** 10.1101/2022.01.26.477883

**Authors:** E. Joe Moran, Maria M. Martignoni, Nicolas Lecomte, Patrick Leighton, Amy Hurford

## Abstract

Empirical observations and mathematical models show that climate warming can lead to the northern (or, more generally, poleward) spread of host species ranges and their corresponding diseases. Here, we explore an unexpected possibility whereby climate warming induces disease spread in the opposite direction to the directional shift in the host species range. To test our hypothesis, we formulate a reaction-diffusion equation model with a Susceptible-Infected (SI) epidemiological structure for two host species, both susceptible to a disease, but spatially isolated due to distinct thermal niches, and where prior to climate warming the disease is endemic in the northern species only. Previous theoretical results show that species’ distributions can lag behind species’ thermal niches when climate warming occurs. As such, we find that climate warming, by shifting both species’ niches forward, may increase the overlap between northern and southern host species ranges, due to the northern species lagging behind its thermal tolerance limit, thus facilitating a southern disease spread. As our model is general, our findings may apply to viral, bacterial, and prion diseases that do not have thermal tolerance limits and are inextricably linked to their hosts’ distributions, such as the spread of rabies from arctic to red foxes.

## Introduction

Many studies have observed shifts in disease range in the same direction as climate warming-induced shifting thermal isoclines (Bellard et al., 2013; Patz et al., 1996; Short et al., 2017). Diseases may spread poleward, or upwards in altitude, when pathogen fitness closely tracks environmental temperature shifts, either due to host or vector responses to temperature (e.g., Lyme disease (Brownstein et al., 2005)), or due to pathogen life stages that are exposed to the environment (e.g., chytrid fungus (Pounds, 2001)). There are many studies investigating the poleward spread of between-host and vector-borne diseases, including malaria (Martens et al., 1995), dengue fever (Hales et al., 2002), bluetongue (Purse et al., 2005), wooly adelgid beetle in hemlocks (Paradis et al., 2008), or beech bark disease (Stephanson and Ribarik Coe, 2017). There are currently no examples of climate-induced disease spread in the opposite direction of climate warming. We hypothesize that when uninfected, susceptible, populations disperse poleward (to where the climate is now warmer), and meet infected populations living at higher latitude, contact between the two populations can facilitate an anti-poleward wave of disease.

Our hypothesis arises as in a multi-host system, where disease can spread to another host species given sufficient between-species contact rates, differences in host ranges can prevent disease transmission, by reducing the contact rates between species. However, climate warming can affect niches geographical extent and, consequently, species distributional area, thereby facilitating disease spread into susceptible populations that have previously been isolated. Indeed, both empirical studies (Menéndez et al., 2006; Talluto et al., 2017) and mathematical models (Hurford et al., 2019; Zhou and Kot, 2011) have shown that in response to climate warming, species may lag behind their shifting thermal tolerance limits. This suggests that for two host species occupying distinct niches along a thermal gradient (for convenience here assumed to be a latitudinal temperature gradient in the Northern Hemisphere) the northern species may lag behind its warm tolerance limit in the South; and as the southern species spreads into the northern limit of its range, the area where the two species overlap increases, thus facilitating disease spread from the northern population to the southern population.

To test whether, and under which conditions, southward disease spread can occur in response to climate warming, we formulate a reaction-diffusion equation model that accounts for disease dynamics for directly transmitted pathogens between spatially structured host populations in a moving habitat. Moving habitat models (Harsch et al., 2014) have been formalized as either reaction diffusion equations (Berestycki et al., 2009; Potapov and Lewis, 2004), or their discrete time analogue: Integrodifference Equations (IDE; Zhou and Kot (2011)), and have been used to study how the speed of climate change impacts population persistence and how population densities respond to shifting habitats. More recently, moving habitat models have been expanded to incorporate infectious agents and species interactions (Leung and Kot, 2015; Kura et al., 2019). Our model incorporates species growth, diffusion, and interaction in a moving-habitat framework, to investigate how the population densities of two host species are affected by a climate-induced shift in the location of their thermal tolerance limits, and how a possible overlap of their expansion ranges may facilitate disease spread.

Our study is not bound to a specific disease, and focuses on systems consisting of two populations with a common pathogen, which are spatially isolated but sufficiently close in space to raise concerns about a possible overlap due to niche shift. We assume that prior to climate warming, there exists an infected northern population, and an uninfected, but susceptible, southern population. The arctic rabies system, for example, lends itself to this formulation of our pre- and post-climate warming scenarios. Indeed, historically, rabies has been endemic in Arctic foxes *(Vulpes lagopus)* (i.e., the “northern population”), while red foxes *(Vulpes vulpes)* (i.e., the “southern population”) have remained disease-free with only sporadic outbreaks (Mörk and Prestrud, 2004; Tabel et al., 1974). The movement of red foxes northward, facilitated by climate change and anthropogenic disturbance, has already led to an increase in overlap among the two species which can be observed in most arctic areas (Gallant et al., 2012, 2020; Savory et al., 2014), and might constitute a threat for potential fast spread of rabies to the south, given the vast distribution of red foxes across Eurasia, North America, part of North Africa and in most of Australia (Hoffmann and Sillero-Zubiri, 2021), with major consequences for human and animal health. Additionally, if rabies is spread southward, rabies’ disease range may overlap with more host species, specifically skunks and raccoons (Finnegan et al., 2002), opening up new transmission pathways. It is therefore imperative to understand how climate warming can contribute to the risk of the southern spread of diseases, for the prevention and management of rabies, as well as other prion and viral diseases.

## Model and Methods

We formulate a temperature-driven moving habitat model based on a reaction-diffusion framework (Cantrell and Cosner, 2004) to understand disease dynamics for directly transmitted pathogens in a warming climate, and in spatially structured host populations. Our model combines disease dynamics with the reproduction, survival, and dispersal of two host populations (i.e., the northern population, characterized by the sub index “n”, and the southern population, with sub index “s”) in a landscape consisting of a thermal gradient, such that each population occupies a distinct region in the North or in the South. We assume that the dynamics characterizing the northern and the southern host populations are identical, except for the thermal tolerance limits of the two populations.

### Spatio-temporal dynamics

Susceptible and infected individuals disperse by random motion, where the dispersal ability is quantified by the diffusion coefficient *D_n_* (for the northern species) and *D_s_* (for the southern species). We assume that susceptible populations exhibit logistic growth, with a temperature-dependent reproductive rate *r_n_(T(x,t))* or *r_s_*(*T*(*x,t*)) (described below, see Eq. (2)), and density-dependent mortality rate *μ_n_* or *μ_s_*. We assume that infectious individuals do not reproduce, and die with a density-dependent mortality rate *ν_n_* or *ν_s_*. Susceptible individuals can become infected by contacting infected individuals in northern or in southern populations alike, where disease transmission occurs at rate *β_nn_, β_ss_, β_ns_* or *β_sn_*, depending on whether the contact has been between two individuals of the same population (northern or southern) or of different populations.

The system of equation describing the spatio-temporal dynamics of the northern and southern populations is given by:

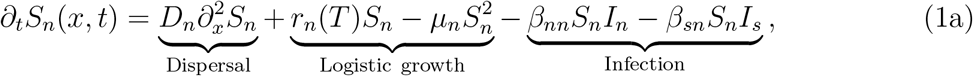

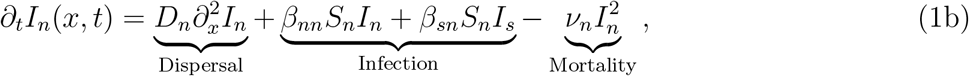

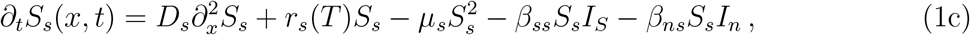

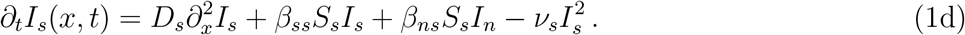

where *S_n_*(*x, t*), *I_n_*(*x, t*), *S_s_*(*x, t*) and *I_s_*(*x, t*) represent the densities of susceptible and infected individuals in the northern and southern populations respectively, at time t and at location x. Although, for application to a specific host-parasite system, the modelling of population growth, disease dynamics, and dispersal may require a more complex framework than that provided in Eq. (1), in order to emphasize the broad validity of our findings we aimed for the simplest possible formulation of the population dynamics, which relies on very minimal assumptions. Possible extensions of the model will be discussed later in this manuscript.

### Temperature, species niches and climate warming

In Eqs. (1a) and (1c), the birth rates *r_n_*(*T*) and *r_s_*(*T*) are represented as functions of temperature *T* = *T*(*x,t*), which depends on the location *x* and on time *t*. Specifically, we assume that birth rates are constant and greater than zero within the species’ thermal tolerance range, identified as the species “thermal” or “fundamental niche”, and zero outside of the thermal tolerance range. We write:

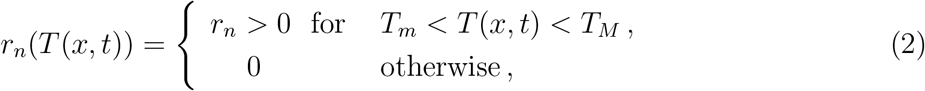

where, *T_m_* and *T_M_* are the lowest and highest temperatures that a species can reproduce at, and an analogous expression can be written for *r_s_* (*T*). While a more gradual change in the net reproduction rate along the temperature gradient may be more realistic (Amarasekare and Savage, 2012), our assumption of a rectangular niche shape represents the least favorable conditions for the northern population to lag behind the southern limit of its thermal tolerance, and therefore, the least favorable conditions for a warming-induced southern wave of infection. Therefore, we expect that if southward disease spread is possible for the rectangular niche shape, this spread will also occur when both species’ niches are assumed to change more continuously as a function of temperature.

Species’ thermal tolerance limits translate into a hospitable region in space (the species niche) where the temperature range is within the indicated limits. We assume the spatial location *x* = [—*L, L*] to be a one-dimensional domain corresponding to a temperature gradient in the Northern Hemisphere, where the temperature decreases gradually from “-*L*” (the “south”) to “*L*” (the “north”) (Fig. 1). We choose *L* = 150 km, and a temperature range prior to climate change extending from 15°C at —*L* to −15° at *L* over 300 km. The default thermal tolerance limits of the northern species are assumed to range from −15°C to −1°C, while the thermal tolerance limits of the southern species range from 1°C to 15°C. The impact of varying those limits will be investigated as described in the next subsection.

**Figure 1:**
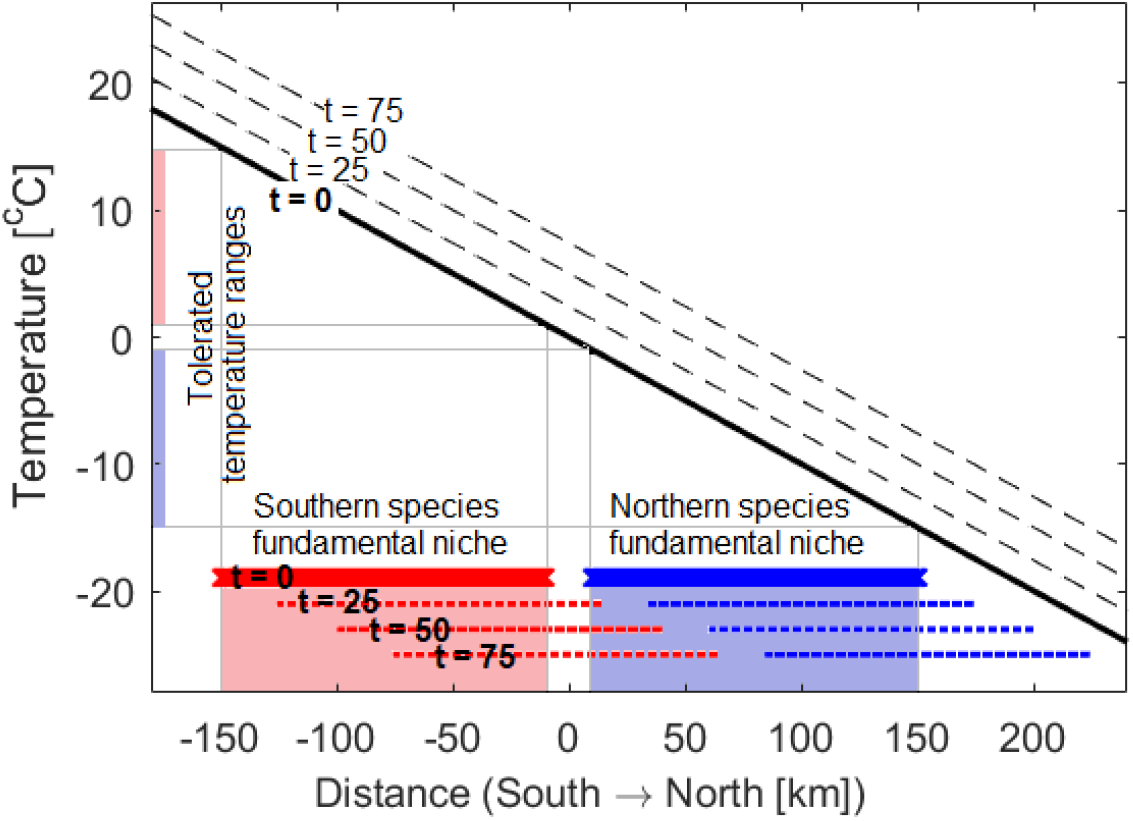
Hypothesized south-to-north temperature gradient as a function of location *x* prior to climate warming (solid black line, *t* = 0) and after 25, 50 and 75 years of climate warming (dashed black lines, *t* = 25, 50, 75). The temperature is assumed to increase 0.1°C per year, which corresponds to a yearly 1 km shift to the north. The location of the fundamental thermal niches of the northern (in blue, color online) and southern (red, color online) populations prior and after climate warming are represented as horizontal solid lines (for *t* = 0) and dashed lines (for *t* = 25, 50, 75). The tolerated temperature ranges of each species prior to climate change are indicated on the vertical axis, where the northern species tolerates lower temperatures (ranging from −15° to −1°), and the southern species tolerates higher temperatures (ranging from 1° to 15°). The impact of varying the Euclidean distance between niches, and thus the thermal tolerance limits of each species, is discussed in Fig. 3b.

Climate warming, beginning at time *t* = 0, causes an increase in temperature by 0.1°C per year (Pachauri et al., 2014), which in our simulations correspond to a northern shift of 1 km per year. Therefore, climate change causes a shift in the thermal tolerance limits (and thus in the fundamental niche) of each species northwards, at a constant rate, and equally at all points in space.

### Simulations

We will focus on the situation where, prior to climate change, species distributions has reached endemic equilibrium, where the disease is present in the northern population only. Although the southern population is also susceptible, it is spatially isolated due to the distinct thermal niche, and thus disease-free. We assume a numerical cutoff value of 0.001, below which population densities are considered to be zero.

As climate change occurs, the temperature gradient is uniformly increased, which results in a spatial shift in the thermal niches of both species. We simulate 75 years of climate warming, and investigate how the time needed for the disease to reach the southern population is affected by the dispersal ability, reproduction, mortality, and disease transmission rate of the two populations, and by variation in the thermal tolerance limits, affecting the Euclidean distance between fundamental niches. Simulations are run in MATLAB R2020b and the computer code is available at https://figshare.com/s/60caec76973c3da640d0.

## Results

Our numerical simulations show that climate warming may induce the southward spread of disease when host species’ ranges shift northwards (Fig. 2). Prior to climate change, the disease reaches endemic equilibrium in the northern population and, because of the spatial isolation arising from the distinct thermal niches, the disease does not spread into the southern population (Fig. 2a). After 25 years of continuous climate warming, the thermal niches of both populations have moved northwards, as have their population densities, but these densities now lag behind their thermal tolerance limits (Fig. 2b). The infected northern individuals (b; blue dashed line) shown south of *x* = 35 km occupy habitat that is too warm, and will ultimately go extinct even if no further climate warming occurs; however, extinction takes time and disease spread to the southern population is enabled via this transient persistence. Indeed, the lag of the northern infected population behind its southern thermal tolerance limit is sufficient to “bridge the gap” to the northern limit of the southern susceptible population (b; right-most red dashed line), allowing the disease to be transmitted to the previously isolated and uninfected southern species. Once disease establishes in the southern population, we observe a wave of infection, which moves southward in space (Fig. 2c and d).

**Figure 2:**
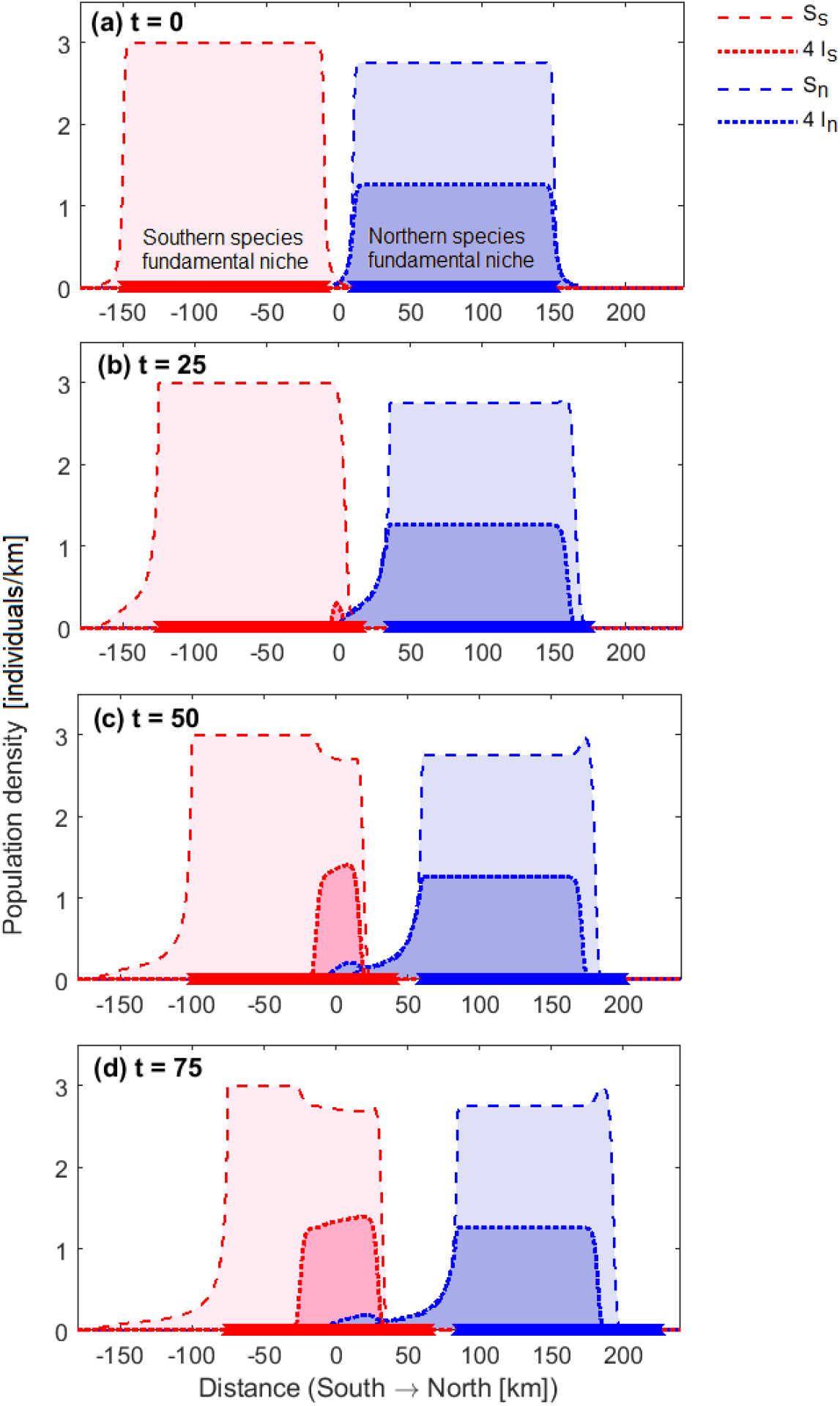
Population dynamics (a) prior to climate change (*t* = 0), and after (b) 25 years, (c) 50 years, and (d) 75 years of climate warming. The southern and northern population densities as a function of the location x are represented in blue and red respectively (color online), with dashed lines representing susceptible individuals and dotted lines representing infectious individuals. Thick blue and red horizontal lines indicate the fundamental niches of the northern and southern species respectively. Prior to climate change, the disease is endemic in the northern population, while the southern population is disease free. Climate change induces a gradual northern shift of both fundamental niches, and disease spread from the northern to the southern population. For visual purposes, the density of the infected populations have been multiplied by 4. Parameters used for the simulation are: *r_n_* = *r_s_* = 1.5, *μ_n_* = *μ_s_* = 0.5, *ν_n_* = *ν_s_* = 3.5, *D_n_* = *D_s_* = 0.3, *β_nn_* = *β_ss_* = *β_ns_* = *β_sn_* = 0.4.

A climate-induced southern spread of the disease is observed only if the thermal tolerance limits, and thus fundamental niches, of the two host species are far enough to be spatially separated before climate warming occurs, but close enough to allow disease spread after climate warming begins. When a southern spread is observed, the Euclidean distance between niches greatly affects the number of years of climate warming needed before disease spread between populations is observed (Fig. 3a). Additionally, southern disease spread requires a high dispersal ability and birth rate of the southern species (Fig. 3b and c), and it is more likely to be observed when the mortality rate of northern infected individuals is low (Fig. 3d). Other model parameters, such as the disease transmission rates and the dispersal ability, mortality, and birth rate of susceptible individuals in the northern population, do not greatly affect the number of years needed till southern spread is observed (see supplementary information, Fig. S.1).

**Figure 3:**
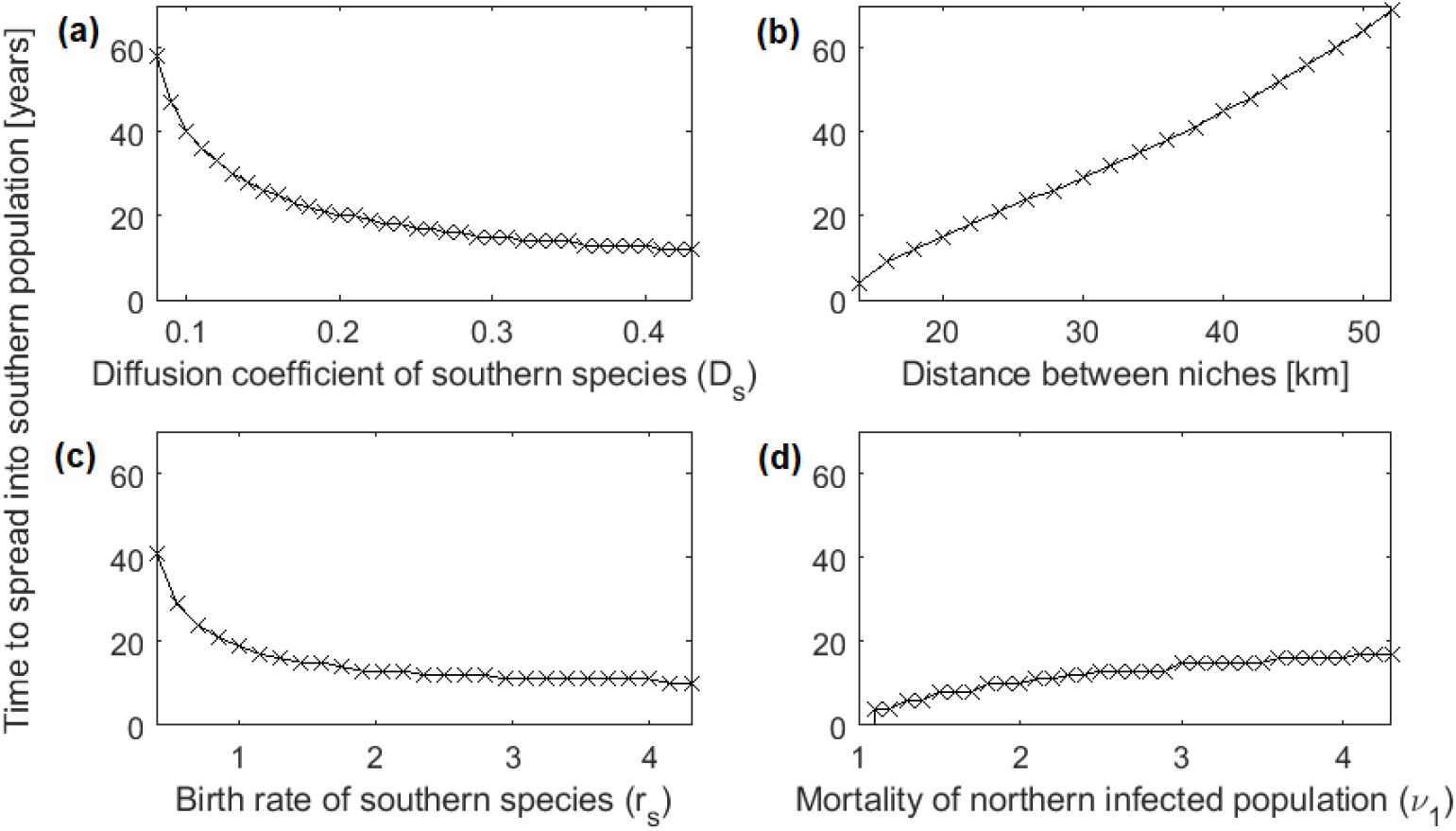
Years of climate warming elapsing before the spread of the disease in a southern population is observed, as a function of (a) the dispersal ability of the southern species (*D_s_*), (b) the thermal tolerance limits of the northern and southern species, determining the Euclidean distance between their fundamental niches prior to climate warming, (c) the birth rate of susceptible individuals of the southern population (*r_s_*), and (d) the mortality rate of the northern infected population (*ν_n_*). Other parameter values are given in Fig. 2.

## Discussion

We find that climate warming can induce the spread of infectious diseases in the direction opposite to host species range shifts. This occurs when lagging species distributions, induced by climate change, connect previously isolated populations, facilitating a southern (or anti-poleward) disease spread. The lag of species distribution behind their thermal tolerance limits has previously been noted for single species moving habitat models (Berestycki et al., 2009; Zhou and Kot, 2011). Specifically, as the niche shifts northwards in response to climate warming, individuals that do not track with their thermal niches do not immediately go extinct in habitat that has recently become inhospitable, but exhibit exponential decay (Amarasekare and Savage, 2012). The area where the population is eventually expected to go extinct has been termed “extinction debt”, and has been demonstrated both empirically (Menendez et al., 2006; Talluto et al., 2017) and theoretically (Hurford et al., 2019; Zhou and Kot, 2011). Here we propose that “extinction debt” areas can facilitate disease spread in the opposite direction of climate warming, via this transient persistence of infected individuals.

We note that the conditions required for the southern spread of disease may be restrictive: 1) there must exist a spatially isolated susceptible, but uninfected population in the south; and 2) the southern population must not be so isolated that individuals cannot disperse into the regions occupied by the lagging infected northern population (made recently suitable for the southern species due to climate warming). High dispersal ability and birth rate of the southern species, as well as a small death rate of infected individuals in the North can also largely determine whether southern spread of disease will be observed, and after how many years of climate warming the spread will occur.

Our counter-intuitive results have implications for epidemic readiness in regions adjacent to areas where disease is endemic. Arctic rabies is an example of a disease system which potentially exhibits the necessary prerequisites for warming-induced southward disease spread. Other host-host disease systems may include arctic fox and raccoon dogs in Europe, which exhibit similar latitudinal distribution and interactions to the arctic – red fox system (Mørk and Prestrud, 2004), or bovine tuberculosis and brucellosis: bacterial pathogens that are endemic diseases in northern bovids, such as the woodland bison of northern Canada (Joly and Messier, 2005; Nishi et al., 2006), and might may spread in southern ungulate populations given climate induced range shifts.

In addition to host-host systems, our model can also apply to host-parasite systems, if a free-living parasite is long-lived and able to withstand warmer temperatures than the host. In such systems, if the pathogen is shed, and the climate later warms, the distribution of the pathogen can lag behind the warm tolerance limit of the host. This can be the case for *Echinococcus multilocularis* for instance, an intestinal parasite endemic to northern latitudes which causes Alveolar echinococcosis disease in carnivorous animals, and can remain infectious in the environment for over 1.5 years (Veit et al., 1995). The presence of multiple hosts of the parasite (such as foxes, wolves, coyotes, or even domestic dogs) raises concerns on whether climate change may contribute to its possible southward movement (Massolo et al., 2014). Also Chronic Wasting Disease (CWD), spread by infectious prions, can persist for more than 2 years in the environment (Miller et al., 2004), and prions from comparable animal diseases (e.g., scrapie disease in sheeps) can persist for up to 16 years (Georgsson et al., 2006).

Our model, despite its simplicity, provides an important first step in raising awareness around the risk of southern disease spread due to climate change. Future modelling efforts should consider different dispersal patterns (Sutherland et al., 2000), different niche structures (Barton et al., 2019), and different temperature dependence of birth and mortality rates (Amarasekare and Savage, 2012; Hurford et al., 2019). The impact that competition between host species might have on reducing the disease transmission risk (Tannerfeldt et al., 2002) should also be quantified. Additionally, host species may experience large year-to-year fluctuations in their population densities (Simon et al., 2019), which may affect disease spread. Models would be needed to show how the spread speed may change under various scenarios, such as higher or lower year-to year variations. Specific parametrization and and/or adaptation of the model to real existing systems, such as those described in the previous paragraphs, can provide useful quantitative insights into when and whether southern disease spread might occur, to support decisions on where to focus disease monitoring efforts.

## Authorship statement

MM and EJM wrote the manuscript. MM wrote the code and completed the analysis, building on earlier code and analysis by EJM. PL, NL and AH motivated the research question and revised the manuscript. AH, MM, and EJM conceived of the analysis.

## Acknowledgement

AH, PL and NL were supported by the National Sciences and Engineering Research Council (NSERC) Discovery Grant (AH: 2014-05413; PL: 2014-03793; NL: 2014-03043). AH and PL were supported by the One Health Modelling Network for Emerging Infections (OMNI). AH was supported by the Emerging Infectious Disease Modelling Consortium. NL was supported by the Canada Research Chair Program, by the Canadian Innovation Fund, and by the Universite de Moncton.

## Supplementary information

**Figure S.1:**
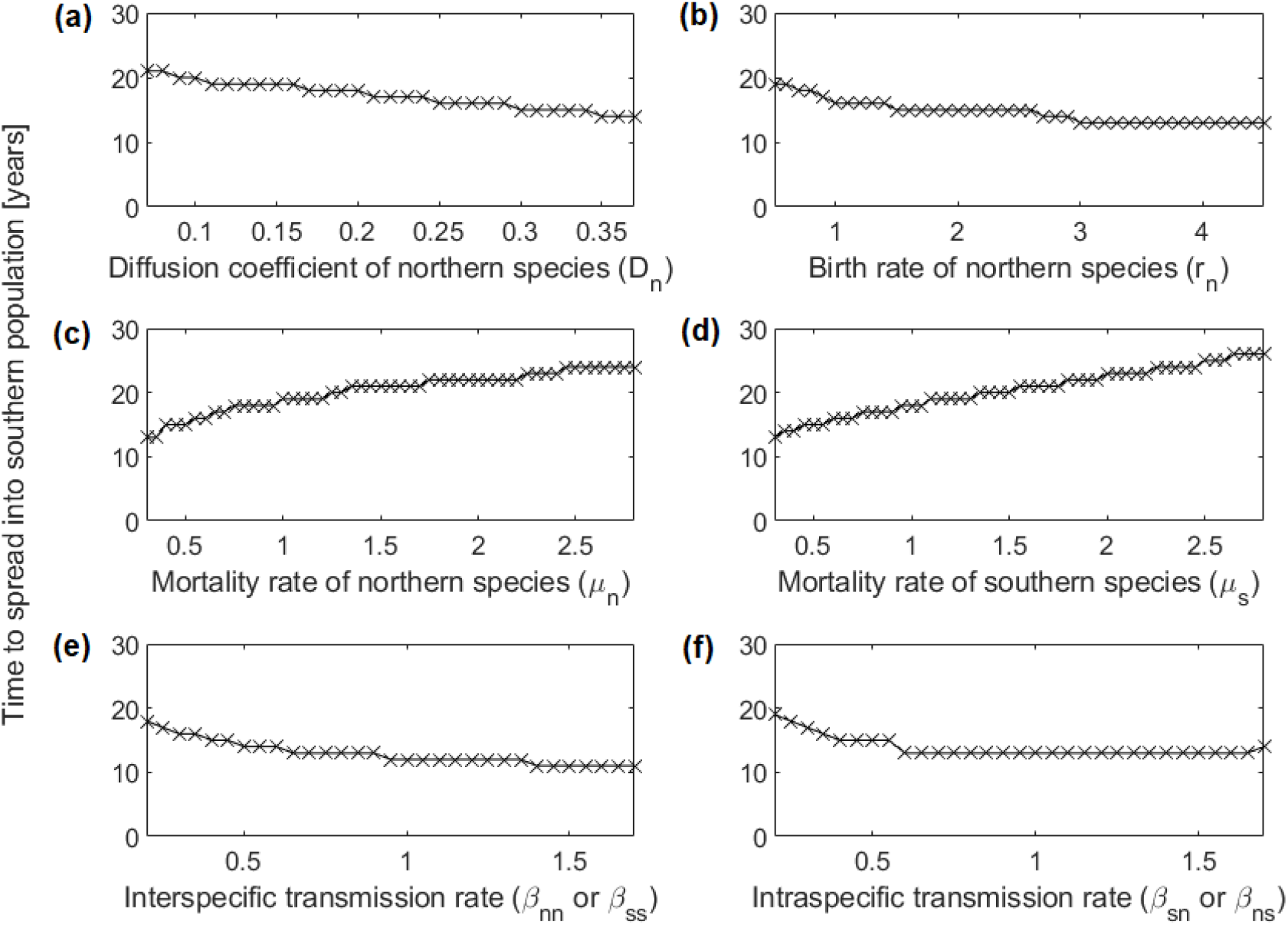
Years of climate warming elapsing before the spread of the disease in a southern population is observed, as a function of (a) the dispersal ability of the northern species (*D_s_*), (b) the birth rate of the northern species (*r_n_*), (c) the mortality rate of the northern species (*μ_n_*), (d) the mortality rate of the southern species (*μ_s_*), (e) the interspecific transmission rate (*β_sn_* and *β_ns_*), (f) the intraspecific transmission rate (*β_nn_* or *β_ss_*). Other parameter values are given in Fig. 2.

